# *Bacillus thuringiensis* bioinsecticides induce developmental defects in non-target *Drosophila melanogaster* larvae

**DOI:** 10.1101/2020.04.30.071563

**Authors:** Marie-Paule Nawrot-Esposito, Aurélie Babin, Matthieu Pasco, Marylène Poirié, Jean-Luc Gatti, Armel Gallet

## Abstract

Bioinsecticides made from the bacterium *Bacillus thuringiensis* (*Bt*) are the best-selling bioinsecticide worldwide. Among *Bt* bioinsecticides, those based on the strain *Bt var. kurstaki* (*Btk*) are widely used in farming to specifically control pest lepidopteran larvae. Although there is much evidence of the lack of acute lethality of *Btk* products for non-target animals, only scarce data are available on their potential non-lethal developmental adverse effects. Using doses that could be reached in the field upon sprayings, we have shown that *Btk* products impair growth and developmental time of the non-target dipteran *Drosophila melanogaster*. These effects are mediated by the synergy between *Btk* bacteria and *Btk* insecticidal toxins, which induces a significant apoptosis of larval enterocytes, resulting in a decreased intestinal capacity to digest proteins. The harmful effects can be mitigated by a protein-rich diet or by adding the probiotic bacterium *Lactobacillus plantarum* into the food. Finally, we showed that the larval midgut maintain its integrity upon *Btk* aggression thanks to both the flattening of surviving enterocytes and the generation of new immature cells arising from the adult midgut precursor cells.

## INTRODUCTION

The use of conventional synthetic chemical pesticides is controversial mainly because of their harmful effects on human health and ecosystems. Public opinion along with governmental policies encourage to reduce their use (Casida and Bryant, 2017; Frisvold, 2018). However, due to the world’s population expansion, the use of pesticides will still be necessary to produce enough food to feed the 9.5 billion humans expected in 2050 (https://population.un.org/wpp/). An emerging solution to reduce the use of conventional pesticides is their replacement with biopesticides.

Among bioinsecticides, *Bacillus thuringiensis* (*Bt*) products are increasingly sprayed in both organic farming and conventional agriculture (Fiuza et al., 2017). *Bt* products are the second most used insecticides in the world with 32 000 tons sold in 2017 (Casida and Bryant, 2017). *Bt* is a Gram-positive spore-forming bacterium belonging to the *Bacillus cereus* group (Vilas-Boas et al., 2007). It was first identified and characterized for its specific entomopathogenic properties due to the presence of specific Cry toxins, which are produced in a crystalline form during the sporulation of bacteria (Rabinovitch et al., 2017). Seventy-eight different strains of *Bt* are currently inventoried (http://www.bgsc.org/), producing a total of more than 300 distinct Cry toxins (http://www.lifesci.sussex.ac.uk/home/Neil_Crickmore/Bt/) having a spectrum of toxicity ranging from nematodes to human tumor cells (Adang et al., 2014; Frankenhuyzen, 2017). However, only four strains are used commercially as bioinsecticides owing to the specific acute toxicity of their Cry toxins to pest larvae: *Bt var. kurstaki (Btk)* and *Bt var. aizawai* to kill lepidopteran larvae, *Bt var. israelensis* to kill mosquito larvae and *Bt var. morrisoni* to kill coleopteran larvae (http://sitem.herts.ac.uk/aeru/bpdb/index.htm). Upon ingestion of commercial formulations made of spores, crystals and additives, Cry protoxins are first released from the crystal before being processed by gut proteases to give rise to the active Cry toxins. Then, these Cry toxins bind to receptors at the surface of the intestinal cells, inducing their death and ultimately the destruction of the gut epithelium. The spores germinate and, thanks to the holes made by Cry toxins in the gut lining, vegetative bacteria invade the internal cavity of the body, causing sepsis which kills the targeted pest in 2 or 3 days. The specificity of the acute Cry toxicity relies on both the capacity of gut proteases to cleave and activate Cry protoxins and on the presence of host receptors specifically recognized by Cry toxins (Adang et al., 2014). Each variant of *Bt* produces between 1 and 6 different Cry toxins. In general, all the Cry toxins contained in a *Bt* variant target a specific phylogenetic order (*e.g.* lepidopteran, coleopteran…) (Palma et al., 2014). The most used *Bt* variant is *Btk*, which produces 5 different Cry toxins (Cry1Aa, Cry1Ab, Cry1Ac, Cry2Aa and Cry2Ab) (Caballero et al., 2020). *Btk* is widely used in forestry and farming to fight lepidopteran larvae.

The increasing environmental dispersion of *Btk* bioinsecticides raises the question of their long-term putative effects on both health and environment. Although there are many data showing that the Cry toxins produced by *Btk* do not display harmful effects against non-target organisms (Haller et al., 2017; Rubio-Infante and Moreno-Fierros, 2016), there are also studies suggesting that Cry toxins and/or *Bt* products may have unintended impacts on non-target animals. For example, the presence of Cry1Ab toxins produced by GM maize in water streams surrounding maize fields delays the growth and increases the mortality of trichopteran species whose larvae have an aquatic lifestyle (Rosi-Marshall et al., 2007; Tank et al., 2010). More recently, Amichot and colleagues have shown that *Btk* bioinsecticides are acutely toxic against *Trichogramma chilonis* females, a tiny egg endoparasitoid hymenopteran used as a biocontrol agent in integrated pest management (Amichot et al., 2016). On vertebrates, Grisolia and colleagues observed that the exposure of *Danio rerio* fry to different *Btk* Cry toxins induced lethality and developmental delay (Grisolia et al., 2009).

Based on these data, we decided to use the model organism *Drosophila melanogaster*, not targeted by *Btk*, to identify and characterize the putative impacts of *Btk* products on the development. While the acute toxicity of the agricultural doses of *Btk* bioinsecticides efficiently kills lepidopteran larva, *Drosophila* larvae, being non-susceptible to these doses, allow the study of non-lethal impacts. In addition, *D. melanogaster* has been used successfully in many studies for deciphering the toxicology of insecticides and is a toolbox widely used to identify disturbed cellular and molecular mechanisms (Scott and Buchon, 2019).

Here, using commercial *Btk* products at two realistic doses, a dose detected on vegetables after one spraying and a dose only tenfold higher (but equivalent to the ten cumulative treatments authorized by the European Union), we observed that *Btk* bioinsecticides induce developmental delay and reduce growth of *D. melanogaster* larvae. We further showed that *Btk* bioinsecticides trigger intestinal cell death and alter protein digestion. Interestingly, we showed that a protein-rich diet or supplementing food with the probiotic *Lactobacillus plantarum* (*L. plantarum*), known to enhance protein uptake (Erkosar et al., 2015), mitigates the impacts of *Btk* bioinsecticide on growth and development. We also found that *Btk* products do not interfere with the commensal flora of the larvae. Finally, we unraveled new cellular mechanisms for maintaining intestinal integrity mounted by the larval midgut epithelium in response to *Btk* aggression, particularly the flattening of enterocytes and the production of new cells from the adult midgut precursors. Overall, our data show that though not lethal for larvae of the non-target organism *Drosophila melanogaster*, agricultural doses of *Btk* bioinsecticides damage gut epithelium and consequently impair larval growth and developmental time.

## RESULTS

### *Btk* bioinsecticide induces developmental delay and growth defect

European and French Food Safety Agencies (Efsa and Anses, respectively) have estimated that the amount of *Btk* spores on fruits and vegetables after treatment can reach 5×10^6^ Colony Forming Units (CFU)/g (ANSES Saisine n° 2013-SA-0039, 2013; EFSA BIOHAZ Panel, 2016). This quantification may be underestimated since up to 10 successive treatments can be applied by farmers (ANSES Saisine n° 2013-SA-0039, 2013) (see also the European directives SANCO/1541/08 – rev. 4 and SANCO/1543/08 – rev. 4). Therefore, we decided to assess the impacts of the ingestion of two *Btk* bioinsecticides (i.e. Delfin and Dipel) on the growth and development of *Drosophila* larvae at two field-realistic doses: 5×10^6^ CFU/g (corresponding to the dose observed after one spraying) and 5×10^7^ CFU/g (a dose which could be reached upon repeated treatments), named hereafter “1X” and “10X”, respectively. We first compared the impacts of Delfin with water (the diluent for commercial products) by contaminating 2g of standard fly medium before depositing 20 eggs aged of 0-4h. We then measured the timing of larval development by performing a pupation curve. In the control experiment (H_2_O), 90% of the larvae pupated at 169h after eggs laying (this threshold will serve as a point of comparison throughout our study, Table 1 summarizes all the data at 10%, 50% and 90% of pupation entry). The 1X and 10X doses of Delfin postponed pupation by 10.5h (+6.2%) and 16h (+9.5%) respectively (Fig. 1A and Table 1). We then assessed the impact of the second bioinsecticide Dipel. Because developmental delay was marked with Delfin at the 10X dose, we used this same dose as a benchmark for Dipel. Strikingly, the pupation delay was increased to reach 25h (+14.8%). Commercial products are made of spores, toxin crystals and additives (see Materials And Methods). Dipel contains 46% of “trade secret” components, while Delfin contains 15% of naphthalene sulfonic acid. To assess the potential involvement of the additives present in Dipel in the augmented delay observed, we plated the commercial product on LB agar, isolated a colony and prepared our own mixture of spores and toxins (the *Btk* strain used in Dipel is named *ABTS-351*). When larvae were fed with the 10X dose of the homemade preparation, we obtained developmental delays similar to those with the Delfin treatments (Fig. 1A and Table 1), suggesting that the unknown additives present in the commercial preparation of Dipel might contribute to the increased delay we observed. Moreover, these data also suggested that the 15% of naphthalene sulfonic acid presents in Delfin had no impact on the developmental timing. Indeed feeding developing larvae with an amount of naphthalene sulfonic acid equivalent to the 10X dose of Delfin had no effect on the developmental timing (Fig. 1B and Table 1). Then, we wondered whether the developmental delay was due to the spores alone, the crystals of toxins alone, or both. First, we fed the developing larvae with the 1X and 10X doses of spores of a *Btk* strain devoid of Cry toxins (*Btk*^*ΔCry*^). Whatever the dose, no developmental delay was observed (Fig. 1B and Table 1). Then, we fed the developing larvae with an equivalent 10X dose of toxin crystals, considering that crystals represent 30% of the dry weight of bioinsecticides (Agaisse and Lereclus, 1995; Monro, 1961; Murty et al., 1994) (Fig. S2). Again, we did not observe any developmental delay (Fig. 1B and Table 1). Therefore, we assumed that the developmental delay we observed was mainly due to the mixture of spores and crystals (Fig. 1A).

**Table 1.**
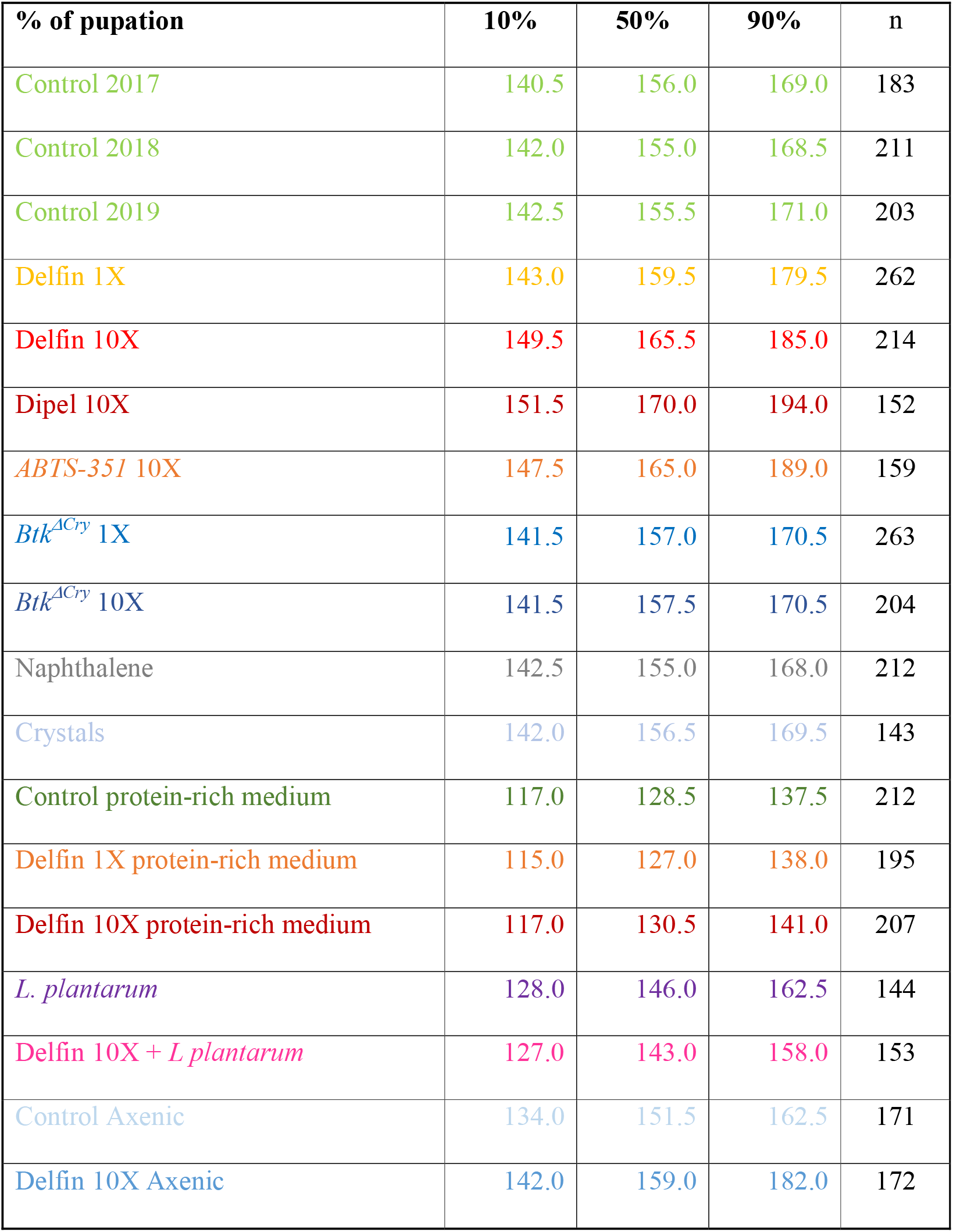
Time in hours when 10%, 50% and 90% larvae enter the pupation stage (immobilized) expressed in hours after egg laying. “n” corresponds to the number of larvae counted in each condition. Note that we compiled the controls per year of experiments (encompassing years from 2017 to 2019).

**Figure 1:**
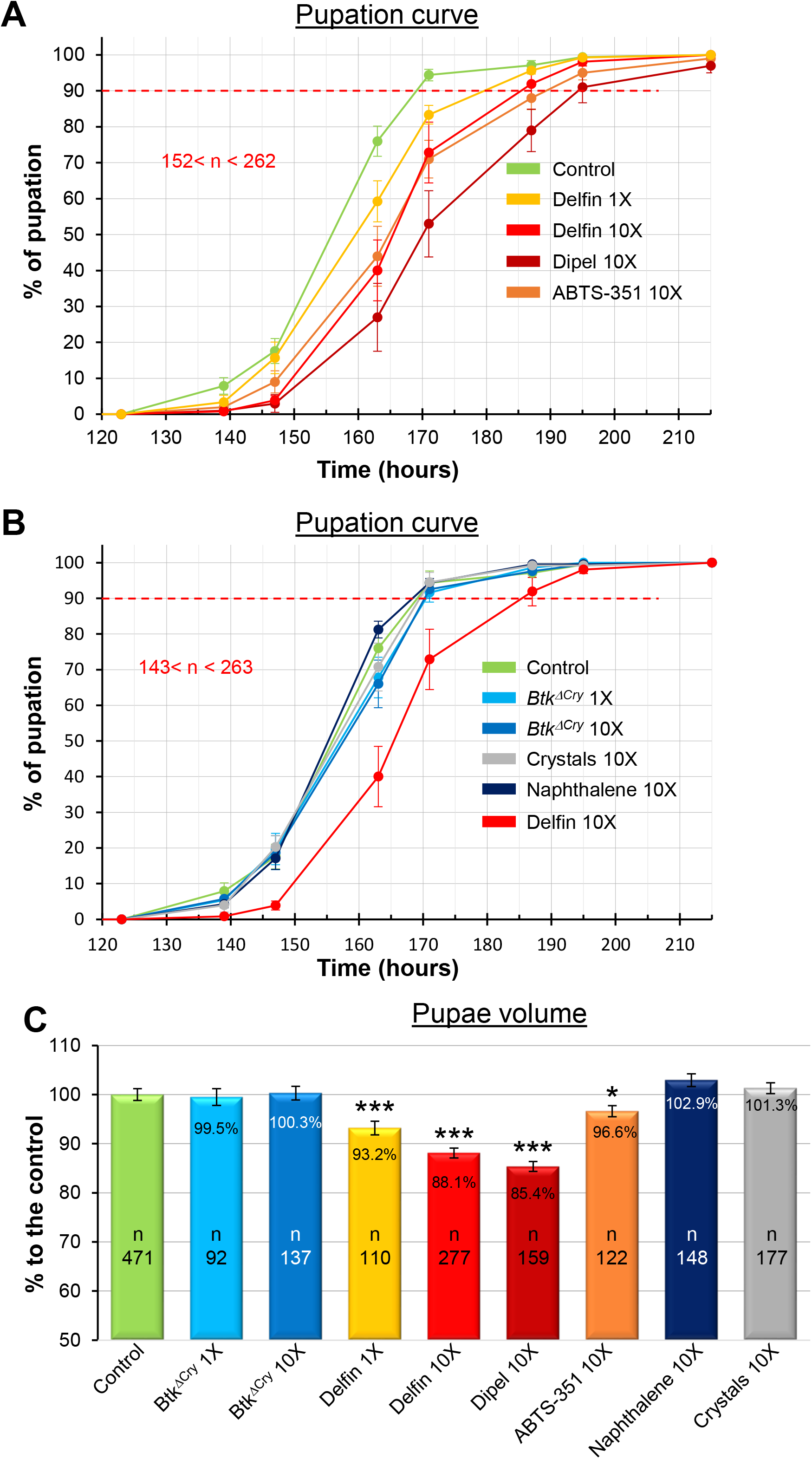
Btk bioinsecticides impacts on developmental delay and growth defects. (**A**) Pupation curve of larvae fed with commercial products (Delfin and Dipel) or with a homemade preparation of spores of the *Btk* strain ABTS-351 (Dipel). In this figure and in all the following figures, the cumulative percentage of pupation is shown over time. (**B**) Pupation curve of larvae fed with a *Btk* strain devoid of Cry toxins (*Btk*^*ΔCry*^) or with Cry toxin containing crystals purified from Dipel or with Naphthalene sulfonic acid at a dose equivalent to the amount found in Delfin 10X. In A and B, red dashed lines mark 90% of the population reaching pupation. (**C**) Pupae volume in the different conditions listed on the abscissa. n= number of individual counted. For the controls, n correspond to the total number of pupae measured when one compiled all the internal controls. Errors bars represent the Standard Deviation of the Mean (SEM). *=P<0.05; ***=P<0.001 compared with controls.

Since growth defects have a well-documented impact on the developmental timing (Hietakangas and Cohen, 2009), we next measured the pupal volume, a known readout to estimate growth defects (Delanoue et al., 2010). Interestingly, we observed a marked growth defect when the larvae were fed with Delfin and Dipel, as well as a weaker one when the larvae were fed with *ABTS-351* (Fig. 1C). We did not detect any growth defect in all the other conditions. Together, our data show that ingestion of food contaminated with a mixture of *Btk* spores and crystals induces developmental delay and growth defect on the model organism *D. melanogaster*, at doses that can be found in treated areas.

Since i) the composition of Delfin is clearly provided by the manufacturer compared to Dipel (see Materials And Methods), ii) Delfin is composed of 85% of spores and toxin crystals while only 54% for Dipel, and iii) the naphthalene sulfonic acid has no effect on the developmental timing and growth, we decided to use only Delfin in the rest of the study.

### *Btk* bioinsecticide does not alter feeding behavior and food intake

The *Btk* load in the midgut could explain the differences observed between *Btk*^*ΔCry*^ and Delfin. When we estimated the amount of *Btk* cells in the midgut of the L3 larvae (Fig. 2A), no difference was observed for 1X or 10X doses of Delfin compared to the equivalent doses of *Btk*^*ΔCry*^. Therefore, the variation in the quantity of *Btk* bacterial cells in the midgut could not explain the disturbed developmental timing and growth observed with Delfin.

**Figure 2:**
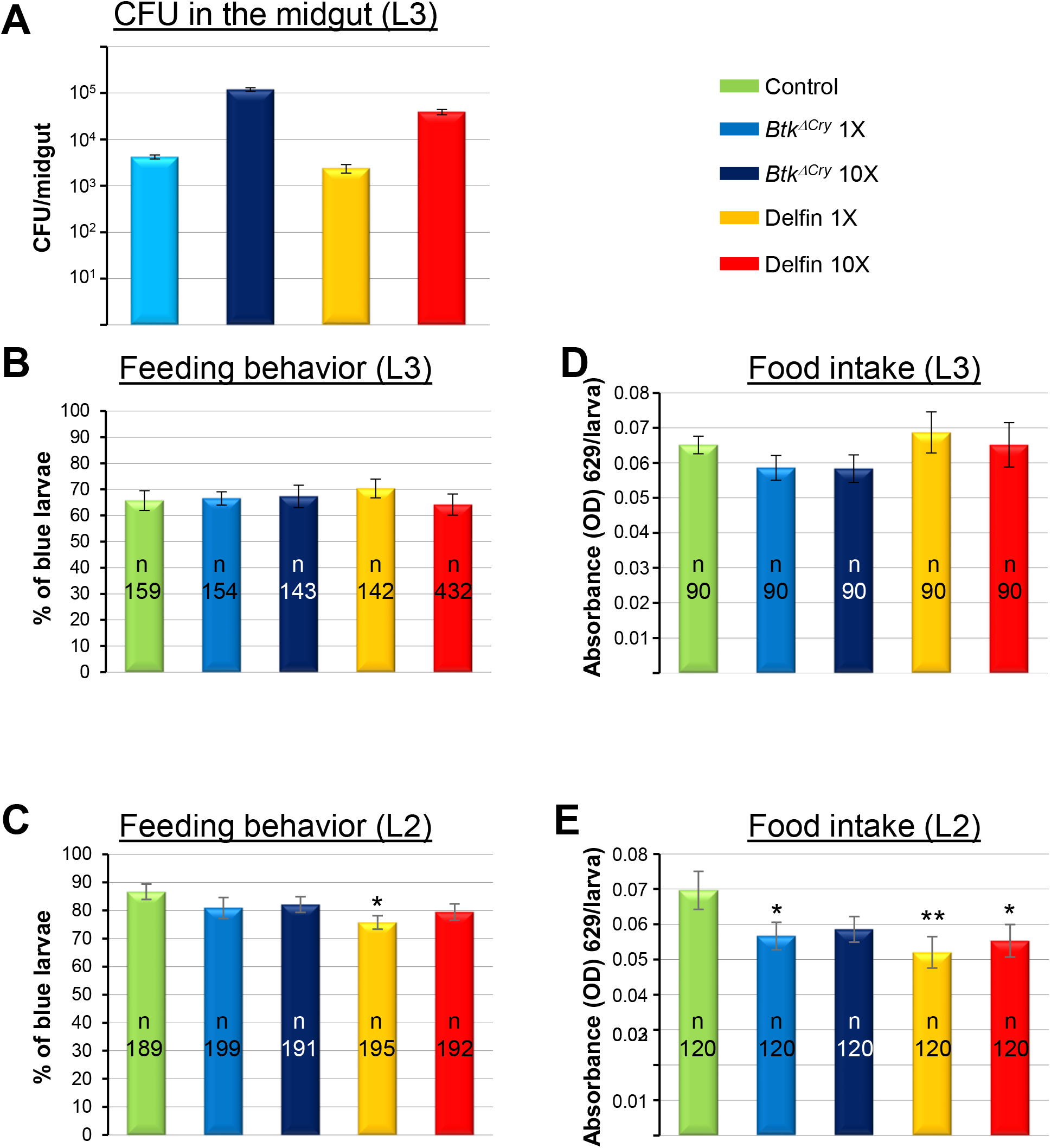
Btk bioinsecticides and larval feeding behavior. (**A**) L3 midgut load of larvae fed with either spores of *Btk*^*ΔCry*^ or Delfin. (**B** and **C**) Percentage of L3 (B) and L2 (C) larvae presenting blue dyed food in the intestine. (**D** and **E**) Amount of dye ingested by L3 (D) and L2 (E) larvae. Green bars: control larvae (fed with water); light blue bars: 1X dose of *Btk*^*ΔCry*^ spores; dark blue bars: 10X dose of *Btk*^*ΔCry*^ spores; yellow bars: 1X dose of Delfin; Red bars: 10X dose of Delfin. n= number of individuals analyzed. Errors bars represent the SEM. *=P<0.05, **=P<0.01 compared with controls.

Adult flies and larvae can sense the presence of harmful microbes, displaying an aversion to the contaminated food (Babin et al., 2014; Keita et al., 2017; Stensmyr et al., 2012; Surendran et al., 2017). Therefore, we tested whether the feeding behavior was affected by the presence of Delfin. We added a blue dye in the food and counted the number of feeding larvae containing blue food in their gut, either in L3 (Fig. 2B) or in L2 stage (Fig. 2C). Although we observed a slight decrease in L2 larvae reared under Delfin 1X dose condition, we did not detect any change in feeding behavior. We then quantified the food intake and, once again, we did not detect any significant modification in the L3 larvae (Fig. 2D), unlike in the L2 larvae, which were affected whatever the condition (excepted for the *Btk*^*ΔCry*^ 10X condition which was not significant) compared to the control (Fig. 2E). Therefore, although food contaminated with *Btk* spores (with or without crystals of toxins) disturbed food intake of L2 larvae but not of L3 larvae, these data do not explain the specific impact of Delfin on the developmental time and growth of the larvae.

### *Btk* bioinsecticide impairs intestinal protein metabolism

Proteins are a key nutrient to drive normal larval development and growth. Increasing protein content in the diet accelerates the developmental time and/or stimulates growth, while an increase in sugar content delays development and a change in lipid amounts has only a marginal impact (Anagnostou et al., 2010; Colombani et al., 2003; Layalle et al., 2008; Reis, 2016; Rodrigues et al., 2015; Soultoukis and Partridge, 2016). Because we have previously shown that the presence of opportunistic bacteria in the adult midgut impede the digestion of proteins (Loudhaief et al., 2017), we wondered whether this could also be the case in larvae in the presence of *Btk* bioinsecticide. While the 10X dose of *Btk*^*ΔCry*^ had no consequence, we observed a decreased capacity of the midgut of L3 larvae to digest proteins in a Delfin dose-dependent manner (Fig. 3A).

**Figure 3:**
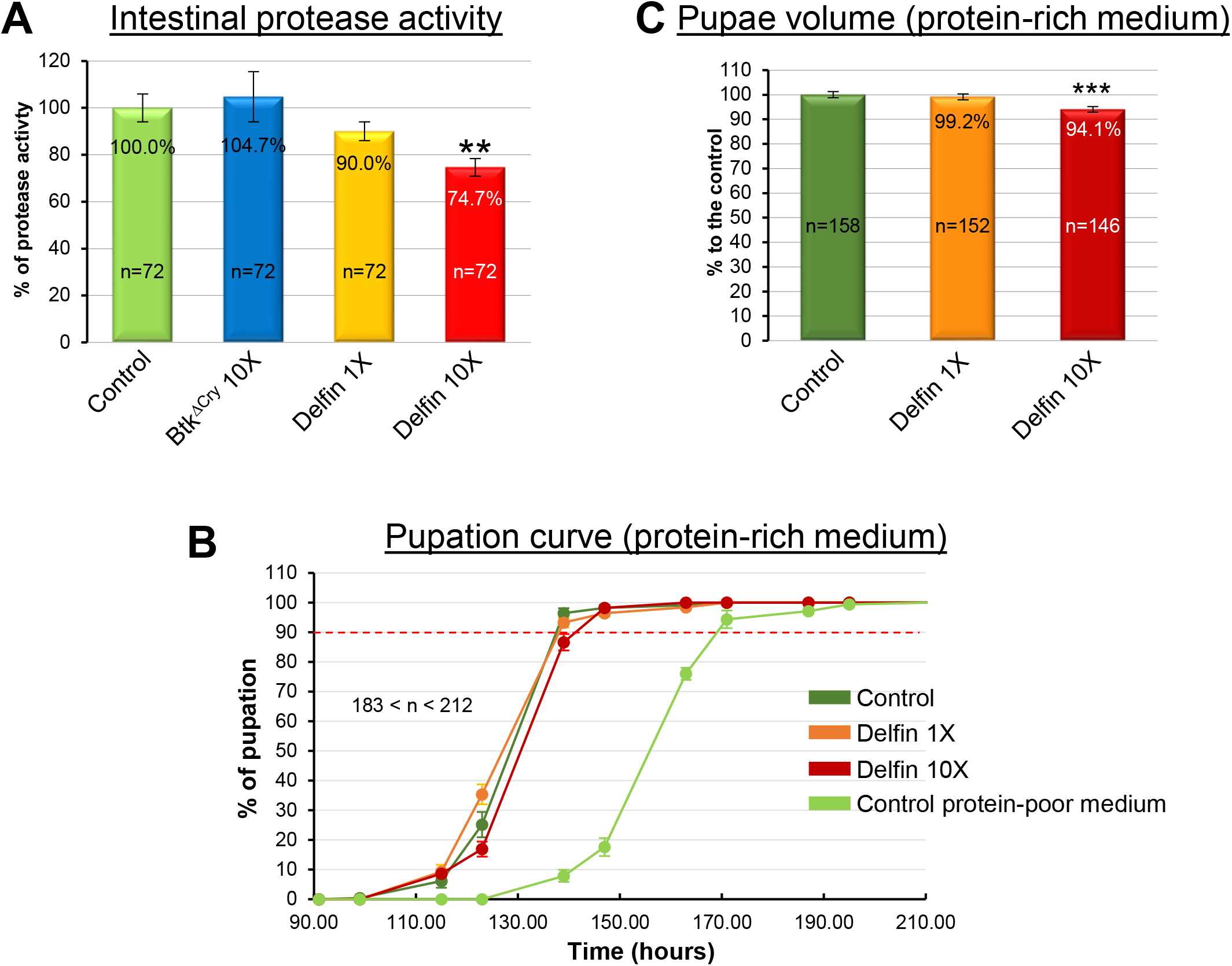
Influence of a protein rich-diet on the Delfin impacts. (**A**) Measurement of the protease activity of the intestine of L3 larvae raised on control medium contaminated with H2O (control) or with 10X doses of spores of *Btk*^*ΔCry*^ or Delfin or 1X dose of spores of Delfin. (**B**) Pupation curve of larvae raised on a protein rich medium (10% yeast instead of 2%) contaminated with 1X or 10X doses of Delfin. Note the shortening of the developmental time of the control larvae raised on the protein-rich medium (10% yeast) compared to larvae raised on the protein poor-medium (2% yeast). (**C**) Pupae volume derived from larvae raised on the protein-rich medium and in the different conditions listed on the abscissa.

We next wondered whether increasing the protein content of the diet could rescue the developmental delay and the growth defects caused by the ingestion of Delfin. Our rearing medium contains 2% yeast as a protein source (see Materials and Methods). We therefore performed a pupation curve with larvae raised on a medium containing 10% yeast with the 1X or 10X doses of Delfin (Fig. 3B). Compared to the control, no difference in the developmental time was observed for the 1X dose of Delfin and only a slight delay for the 10X dose (Table 1). Noteworthy, the developmental time for all conditions on the protein-rich medium was shortened compared to the control on our conventional medium (light green curve in Fig. 3B and Table 1). Similarly, the pupal volume (Fig. 3C) was completely restored for the 1X dose of Delfin and partially rescued for the 10X dose of Delfin (5.9% volume loss instead of 11.9% in the poor-protein diet). Taken together, our data suggest that Delfin impairs protein digestion, and thus gut functions, which can be rescued by adding extra dietary protein.

### The commensal bacterium *L. plantarum* helps to overcome *Btk* bioinsecticide impacts

It has been recently shown that the commensal bacterium *L. plantarum* promotes the juvenile systemic growth of *Drosophila* larvae and infant mice by stimulating digestion and absorption of proteins, and protects the child against sepsis upon undernutrition conditions (Erkosar et al., 2015; Panigrahi et al., 2017; Schwarzer et al., 2016; Storelli et al., 2011; Tefit and Leulier, 2017). Therefore, we wondered whether complementing our conventional medium with *L. plantarum* would compensate for the impacts of the 10X dose of Delfin. First, we observed that adding *L. plantarum* in the conventional medium shortened the developmental time of control larvae (Fig. 4A, purple curve compared to green curve and Table 1). Interestingly, the presence of *L. plantarum* fully counteracted Delfin-dependent developmental delay (Fig. 4, pink curve compared to red curve and Table 1). We next observed that the presence of *L plantarum* in the conventional medium reduced the pupal size of the control larvae by 4.4% compared to the control without *L. plantarum* (Fig. 4B, purple bar compared to green bar) but fully rescued the growth defects induced by the 10X dose of Delfin (Fig. 4B pink bar). Since ingestion of *L. plantarum* and its establishment in the midgut (Loudhaief et al., 2017) might compete for the presence of *Btk* and not act on protein intake as expected, we also controlled the number of *Btk* CFUs in the absence or presence of *L. plantarum* in the midgut of L3 larvae and we found no difference (Fig. 4C). Taken together, our data demonstrated that stimulation digestion and/or absorption of proteins by the commensal bacterium *L. plantarum* helps to overcome the adverse effects of the *Btk* bioinsecticides on larval development.

**Figure 4:**
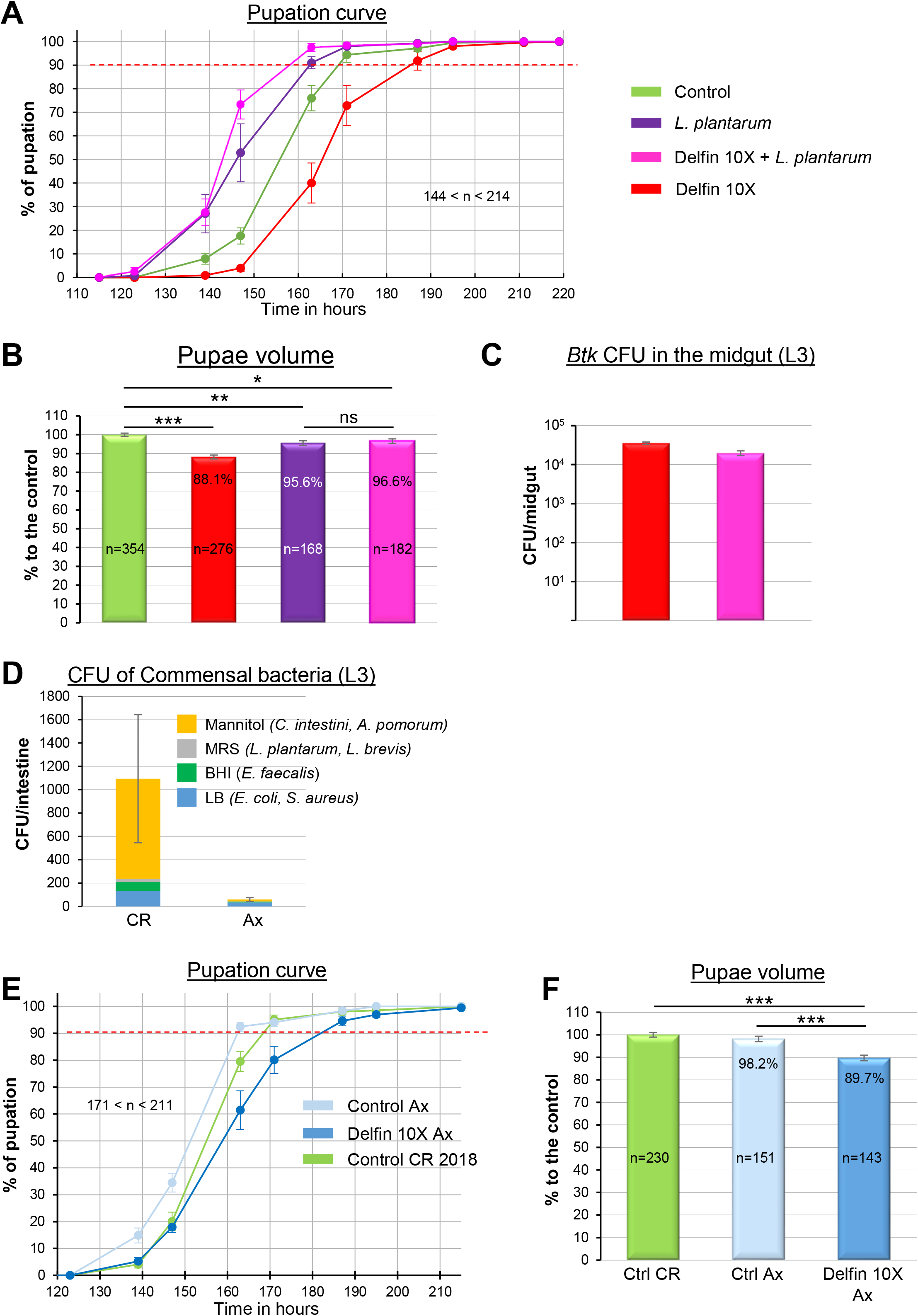
*Btk* bioinsecticides interaction with commensal flora. (**A**) Pupation curve of larvae reared on a medium complemented with 10^6^ CFU of *L plantarum* and contaminated with Delfin 10X (pink) or not (purple) compared to control larvae (green) or Delfin 10X only (red). (**B**) Pupae volume derived from larvae raised on a medium complemented with 10^6^ CFU of *L plantarum* and contaminated with Delfin 10X (pink) or not (purple) compared to control pupae reared on control medium (green) or medium contaminated with Delfin 10X only (red). (**C**) *Btk* load in the midgut of L3 larvae reared on a medium contaminated with Delfin 10X and complemented (pink) or not (red) with 10^6^ CFU of *L plantarum*. (**D**) Commensal bacterial load in the midgut of L3 larvae of our *Drosophila* Canton S strain reared on our conventional medium (left bar) or reared in axenic condition (right bar). The *Commensalibacter intestini* and *Acetobacter pomorum* grow on mannitol plate (orange), *L plantarum* and *L brevis* grow on MRS plate (grey), *Enterococcus faecalis* grows on BHI plate (green) and *Escherichia coli and Staphylococcus aureus* grow on LB plate (blue). (**E**) Pupation curve of axenic larvae raised on a medium contaminated with Delfin 10X (dark blue) or not (light blue, control Ax) compared to larvae conventionally reared (CR, green). (**F**) Pupae volume derived from conventionally reared larvae (green) or from axenic larvae raised on a sterile medium contaminated with Delfin 10X (dark blue) or not (light blue).

### The impacts of *Btk* bioinsecticide do not rely on commensal bacteria disturbance

*L. plantarum* is a widespread commensal bacterium of the *Drosophila* midgut (Broderick and Lemaitre, 2012; Chandler et al., 2011; Fast et al., 2018). Because complementing the larval diet with *L. plantarum* rescued the developmental delay and growth defects induced by *Btk*, we wondered whether the indigenous commensal flora of larvae reared on our conventional medium (i.e. poor in protein) was inhabited by *L. plantarum*. Only few *L. plantarum* and/or *L. brevis* (belonging to the family of *Lactobacillaceae*) were present in the commensal flora of our larvae (Fig. 4D). The dominant bacteria family we identified belonged to *Acetobacteraceae* known to be common in flies feeding on sugary, acidic and alcoholic food (Chandler et al., 2011; Téfit et al., 2018). This observation was in agreement with the composition of our conventional rearing medium poor in protein but quite rich in simple sugar (i.e. sucrose, see Materials And Methods) and it suggested that the *Btk* effects we observed were independent of a putative deleterious impact on the indigenous *L. plantarum* of our larvae.

However, we could not rule out an impact of the *Btk* bioinsecticide on other members of the commensal flora. To test this hypothesis, we generated an axenic fly strain (see Materials And Methods). Axenic larvae (Fig. 4D) were further fed on our conventional medium treated or not with the 10X dose of Delfin. The developmental time of the control axenic larvae was slightly shorter than those of the conventionally reared (CR) larvae (i.e. non-axenic) (Fig. 4E and Table 1). Nevertheless, axenic larvae infected with Delfin 10X displayed a 19h developmental delay compared to the control axenic larvae (+11.7%) (Fig. 4E and Table 1), which was similar to the delay observed with *Btk*-treated CR larvae (see Fig. 1). Then, while axenic larvae gave rise to pupae with a similar volume to that of CR larvae (Fig. 4F), Delfin 10X still induced a reduction of pupal volume in axenic larvae compared to the axenic control (Fig. 4F) in a magnitude similar to what we observed for Delfin 10X with CR larvae (see Fig. 1). Taken together our data strongly suggest that Delfin ingestion affects developmental time and growth independently of the commensal bacteria.

### *Btk* bioinsecticide induces midgut perturbations

Because we had previously shown that the ingestion of *Bt*k vegetative cells by adult *Drosophila* reduced the accumulation of lipids in enterocytes (Loudhaief et al., 2017), we wondered whether the commercial *Btk* product did the same in larvae. In unchallenged L3 larvae, lipid droplets strongly accumulate in enterocytes in the most anterior part of the midgut and weakly in the posterior midgut (Fig. 5A) (Palm et al., 2012). Instead of lipid droplets depletion, feeding the larvae with Delfin 10X provoked an accumulation of lipid droplets in the posterior larval midgut (Fig. 5E). Furthermore, lipid droplets accumulated apically, facing the lumen (Fig. 5F and F’ compared to 5B and B’). Such an accumulation was not observed when developing larvae were fed with *Btk*^*ΔCry*^ spores (Fig. 5C, D and D’). Interestingly, it has been recently demonstrated that the pore-forming toxins hemolysin and monalysin produced respectively by the enteropathogenic bacteria *Serratia marcescens* (*Sm*) and *Pseudomonas entomophila* (*Pe*) can induce an apical accumulation of lipid droplets in adult enterocytes of *Drosophila*, honeybees and mice as well as in epithelial Caco-2 cells in culture. This phenomenon precedes the expulsion of cytoplasm required to get rid of toxins from the enterocytes (Lee et al., 2016). Since the Cry toxins produced by *Btk* are also pore-forming toxins (Adang et al., 2014), we fed developing larvae with purified toxin crystals and checked for the accumulation of lipid droplets. Unexpectedly, we did not observe any accumulation of lipid droplets (Fig. 5G, H and H’). However, feeding our developing larvae with the naphthalene sulfonic acid (the additive present in the commercial products, see Supplemental Materials) provoked an accumulation of apical lipid droplets posterior to the acidic domain (Fig. 5I, J and J’). To further assess whether such a lipid accumulation in the midgut would impair lipid metabolism in the whole larva, we quantified the circulating lipids (the diacylglycerides - DAG) in the hemolymph and the stored lipids (triacylglycerides - TAG) (Tennessen et al., 2014) in mid-third instar larvae. We did not notice any significant perturbation in lipid metabolism (Fig. 5K) though a downward trend was visible for circulating DAG in larvae fed with Delfin 10X. Thereby, it is possible that such an apical accumulation of lipid droplets serves for the trap and elimination of the naphthalene sulfonic acid similarly to what happens for the elimination of the *Sm* and *Pe* pore-forming toxins (Lee et al., 2016).

**Figure 5:**
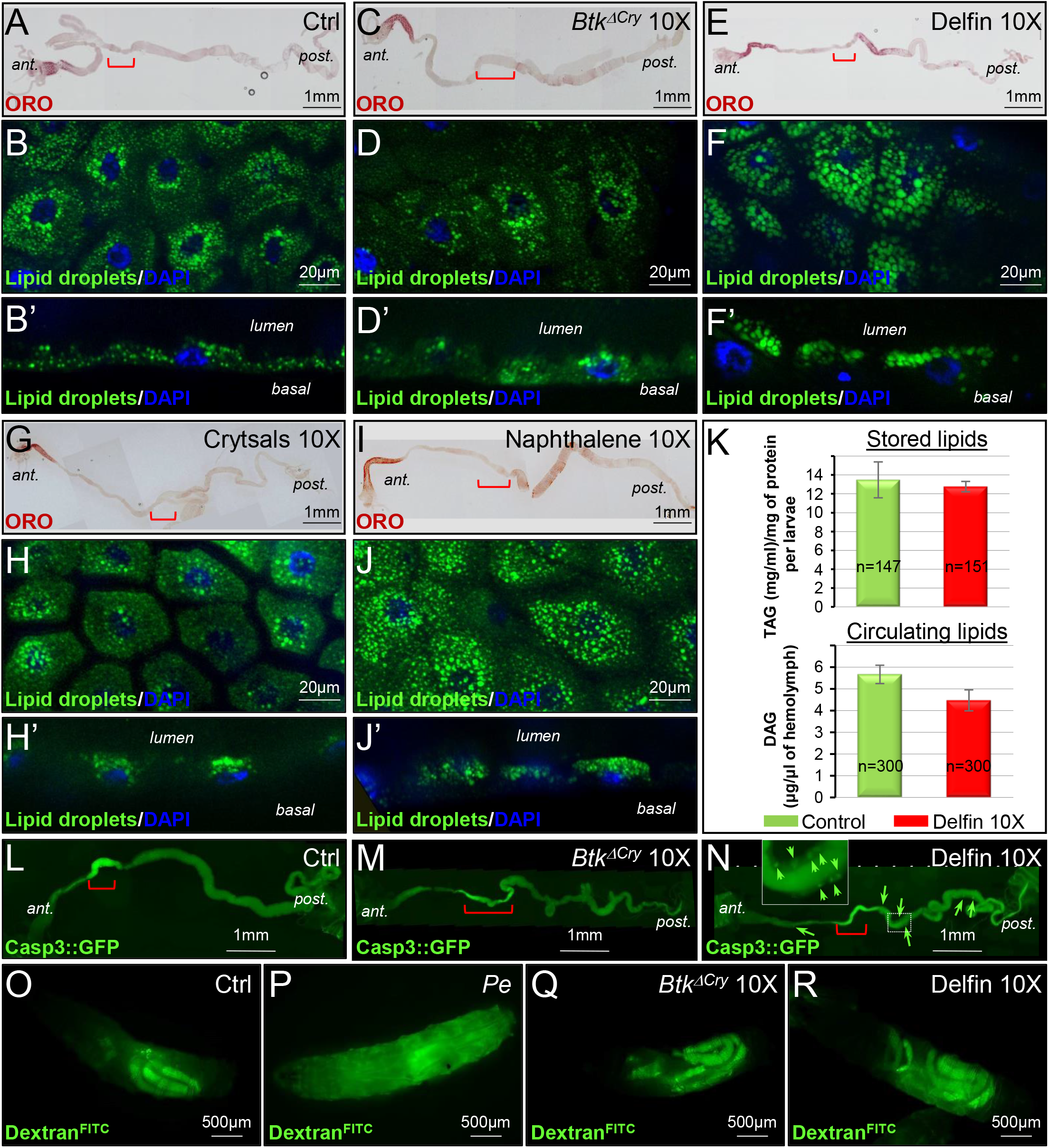
*Btk* bioinsecticides impacts on gut integrity. L3 larvae raised on a control medium (**A-B’**) or a medium contaminated with *Btk*^*ΔCry*^ 10X (**C-D’**), Delfin 10X (**E-F**), purified Crystals 10X (**G-H’**), naphthalene 10X (**I-J’**). (**A-J’**) Labelling of lipid droplets with ORO (red, A, C, E, G and I) or with Bodipy (green, B, B’, D, D’, F, F’, H, H’, J and J’). DAPI staining in blue marks the nuclei. Midguts are oriented anterior to the left. Red brackets delineate the acidic region of the midgut. (B’,D’, F’, H’ and J’) are x/z section with the apical pole of the epithelium facing the lumen. (**K**) Stored (TAG, upper panel) and circulating (DAG, lower panel) lipid dosages in control larvae (green bars) and in larvae raised on Delfin 10X contaminated medium (red bars). (**L-N**) *myo1A>Casp::GFP* larvae raised on control a medium (L) or on a medium contaminated with *Btk*^*ΔCry*^ 10X (M) or Delfin 10X (N). Arrows point enterocytes in which Caspase 3 is activated. (**O-P**) L3 larvae raised on a medium complemented with Dextran^FITC^ (O) or with Dextran^FITC^ and contaminated with *P. entomophila* (P), with *Btk*^*ΔCry*^ 10X (Q) or Delfin 10X (R).

In adult *Drosophila*, pathogenic and opportunistic bacteria can induce enterocyte apoptosis upon ingestion (Apidianakis et al., 2009; Loudhaief et al., 2017). Therefore, we verified the appearance of dying cells in the midgut using the Caspase3::GFP sensor to visualize apoptosis in living tissue (Schott et al., 2017) (see Materials And Methods). While unchallenged and *Btk*^*ΔCry*^ fed larvae did not display any obvious apoptosis, the developing larvae fed with Delfin harbored apoptotic enterocytes mainly in the posterior part of the intestine (Fig. 5L-M and Fig. S3). Because in target lepidopteran larvae, Cry toxins make holes in the gut lining, breaking down the gut barrier and allowing *Btk* cells to penetrate inside the internal milieu (Adang et al., 2014), we checked whether the sealing of the gut epithelium was maintained even in presence of widespread apoptosis. We fed our developing larvae with the small fluorescent molecule Dextran-FITC that is unable to cross the midgut epithelium (Fig. 5O) unless the intestinal barrier is broken, allowing the fluorescence to spread throughout the larva like in the case of *Pe* infection (Fig. 5P). Interestingly, *Btk*^*ΔCry*^ spores as well as Delfin did not induce the loss of gut barrier function (Fig. 5Q and R). Thus, our data show that *Btk* bioinsecticide hampers intestinal lipid homeostasis and induces enterocyte apoptosis without causing the rupture of the intestinal barrier. Such physiological perturbations may be responsible for the developmental delay and the growth defect observed.

### Enterocyte flattening and incomplete differentiation of Adult Midgut Precursors help to maintain intestinal integrity

Intestinal physiology can also be impeded by a disturbance of the apico-basal polarity of the gut epithelium. In the control L3 larval midgut, Armadillo (Arm, β-Catenin), a component of the adherens junction, marks the basolateral compartment of the enterocytes and Disc Large (Dlg), a component of the septate junction, marks the sub-apical compartment. Phalloidin that binds to F-Actin reveals both the apical brush border (facing the lumen) and the basal visceral muscles that surround the midgut lining (Fig. 6A and A’). This organization is similar to the adult midgut (Chen et al., 2018; Loudhaief et al., 2017). Moreover, Arm also delineates the islets of the <u>a</u>dult <u>m</u>idgut <u>p</u>recursors (AMPs), which will give birth to the adult midgut during metamorphosis (Fig. 6A) (Jiang and Edgar, 2009; Mathur et al., 2010; Micchelli et al., 2011). When larvae were fed with the 10X dose of *Btk*^*ΔCry*^ spores, we did not observe any major perturbations of the architecture of late L3 midguts (Fig. 6B and B’). However, we observed an increased number of AMPs per islet (Fig. 6D) correlated with an increase in AMP division (Fig. 6E). When larvae were fed with Delfin 10X, we also observed an increased number of AMPs by islet (Fig. 6D) and a huge increase in AMP proliferation (Fig. 6E) though at the expense of the number of islets (Fig. 6F). Moreover, we also observed an increase in the enterocyte surface (Fig. 6C and G). Interestingly, such an increase in enterocyte surface did not rely on a higher ploidy (Fig. 6J) but rather on a change in cell shape, i.e. enterocyte flattening (Fig. 6C’ and H). Finally, we noticed the appearance of uncharacterized cells having an “intermediate” size. These cells which were intercalated between large enterocytes and the AMP islets, were neither mature polyploid enterocytes nor diploid AMPs or enteroendocrine cells (Fig. 6I and I’). This observation was confirmed by analyzing the ploidy of the nuclei. In the control larvae, 89% of the polyploid cells underwent 4 endocycles of replication (32C nuclei, C corresponding to chromatin copy number) and 10% reached 5 cycles of endoreplication (64C) (Fig. 6J). When larvae were raised on medium contaminated with *Btk*^*ΔCry*^ spores, 74% corresponded to 32C nuclei and 25% to 64C nuclei (Fig. 6J). When larvae were fed with Delfin 10X, a new population of cells with 16C, 8C and 4C nuclei appeared, representing 37% of the population of polyploid cells (Fig. 6J). While the adult midgut quickly regenerates upon an oral infection thanks to the proliferation of adult intestinal stem cells and the differentiation of their daughter cells (the enteroblasts) into new enterocytes which repair the damages (Bonfini et al., 2016), such a mechanism has never been observed in *Drosophila* larvae. Indeed, the number of enterocytes composing the L3 intestine is defined during embryogenesis (Miguel-Aliaga et al., 2018) and the larval gut grows without cell division but by increasing the ploidy and consequently the size of enterocytes. However, our data suggested that upon Delfin ingestion, larval gut homeostasis is perturbed and the apparition of undefined cells is promoted. In the larval intestine, AMPs are the only cells capable of division (Jiang and Edgar, 2009; Mathur et al., 2010; Micchelli et al., 2011). Therefore, we hypothesized that the unknown cells having a nucleus ranging from 4C to 16C (Fig. 6J) and adjacent to the AMP islets (Fig. 6I) could arise from the differentiation of those AMPs. To test our hypothesis, we used the lineage tracing genetic tools developed in *Drosophila* named ReDDM (Antonello et al., 2015). The principle of this lineage tracing is based on the expression of both a labile GFP and a stable RFP within the AMPs thanks to the use of a promotor specifically expressed in AMPs and inducible at will (see Materials And Methods). Accordingly, AMPs express both GFP and RFP, while the progenies, which are not AMPs, only express RFP. In control larvae, GFP and RFP were only present in the islet cells (Fig. 6K) (Jiang and Edgar, 2009; Mathur et al., 2010; Micchelli et al., 2011). When larvae were fed with *Btk*^*ΔCry*^ spores, GFP and RFP were also only expressed in the same AMP cells (Fig. 6L). In contrast, when larvae were fed with Delfin 10X, cells harboring only RFP were now detectable. Furthermore, the newborn cells had nuclei of intermediate size ranging from 4C to 16C (Fig. 6M). This result demonstrated that upon aggression by *Btk* bioinsecticides, the larval midgut is able to produce new cells, most probably immature enterocytes arising from AMPs. Altogether, our data suggest that the larval intestine maintains its integrity upon aggression through two mechanisms, i) enterocyte flattening which increases the cell surface without new cycles of endoreplication and ii) producing new immature enterocytes. Both mechanisms may allow the intestine to maintain the intestinal barrier, avoiding the bacteria to penetrate into the internal milieu and therefore preventing septicemia.

**Figure 6:**
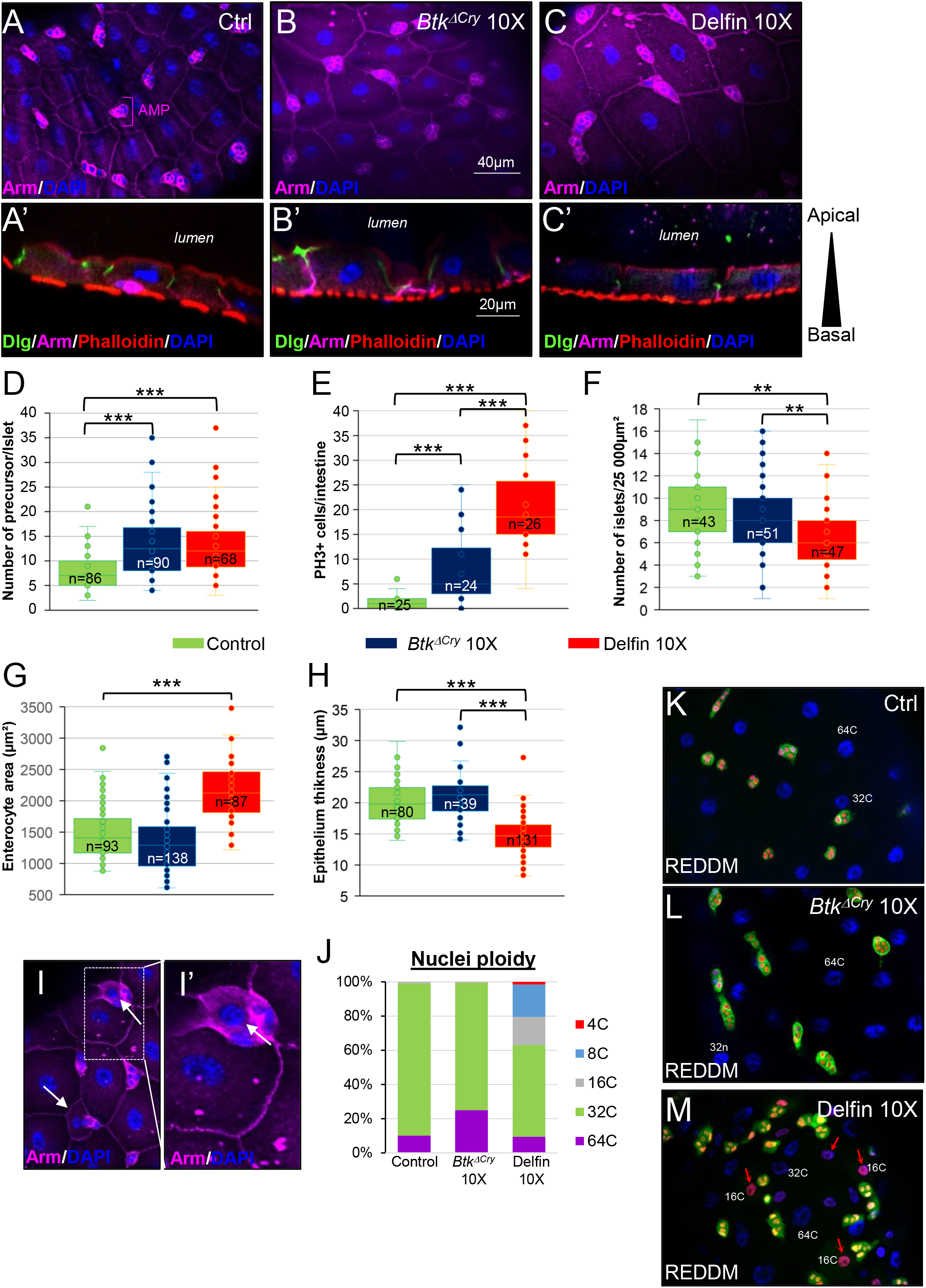
Larval midgut regeneration upon aggression. (**A-C’**) Immunolabelling of *dlg::GFP* L3 midgut from larvae raised on control medium (A and A’), on medium contaminated with *Btk*^*ΔCry*^ 10X (B and B’) or Delfin 10X (C and C’). Midguts were stained with the adherens junction marker Armadillo (Arm, purple) that strongly delineates the Adult Midgut Precursors (AMP) (A, B and C) and marks the basolateral compartment of the enterocytes (A’, B’ and C’). DAPI (blue) marks the nuclei and Phalloidin (red) labels the visceral muscle surrounding the intestine and the apical pole of the enterocytes facing the lumen (in A’, B’ and C’). Dlg::GFP (green) marks the sub apical compartment of the enterocytes. (**D-H**) L3 midgut from larvae raised on control medium (green bars), on medium contaminated with *Btk*^*ΔCry*^ 10X (dark blue bars) or Delfin 10X (red bars). (**D**) Number of AMPs per islets, (**E**) number of dividing AMPs per intestine (mitotic AMPs are labeled by the Phospho-Histone 3 (PH3) immunostaining), (**F**) Number of AMP islets (an islet contains more than one diploid cell) per unit of surface, (**G**) enterocyte area and (**H**) epithelium thickness in the posterior midgut (3 independent measures were taken by image capture). (**I** and enlargement in **I’**) *dlg::GFP* L3 midgut from larvae raised on medium contaminated with Delfin 10X and stained for Arm (purple) and DAPI (blue). Arrows point the cells with intermediate size (smallest than enterocytes but biggest than AMPs). (**J**) Assessment of enterocyte ploidy in posterior L3 midgut from larvae raised on control medium, on medium contaminated with *Btk*^*ΔCry*^ 10X or Delfin 10X (C corresponding to chromatin copy number). (**K-M**) Lineage tracing of AMP progeny in the *w; esg-Gal4, UAS-GFP/+; UAS-H2B::RFP, tub-Gal80*^*ts*^/+ (esg REDDM) L3 midgut from larvae raised on control medium (K), on medium contaminated with *Btk*^*ΔCry*^ 10X (L) or Delfin 10X (M). DAPI (blue) marks the nuclei. **=P<0.01, ***=P<0.001 compared with controls.

## Discussion

The long-term effects of chronic exposure to low doses of potential noxious products cannot be easily revealed by conventional epidemiological studies (which are long lasting and expensive) or conventional (eco)toxicological approaches based on acute toxicity. The use of amenable and cheap laboratory models becomes of prime importance to pinpoint discrete chronic adverse effects. *D. melanogaster* fulfills all the required criteria to be such a model. Indeed, *Drosophila* is easily handling, cheap to rear and the availability of genetic, cellular and molecular tools allows to detect tenuous effects and to decipher the underlying physiological mechanisms impaired (Scott and Buchon, 2019). Moreover, the main physiological functions and genetic networks governing the *Drosophila* gut physiology are well conserved among the animal kingdom (Pasco et al., 2015), allowing to provide informative outputs for both human health and wildlife.

*Btk* bioinsecticides have recently been shown to induce larval lethality and delay the imago emergence of many different *Drosophila* species at doses ranging from 5×10^7^ to 1×10^9^ CFU of *Btk* spores/g of fly food medium (Babin et al., 2019; Cossentine et al., 2016). Larval lethality is observed for the highest doses (Babin et al., 2019) and could be explained by the cross-order toxicity of the Cry2A toxins (Cry2Aa and Cry2Ab are present in *Btk* bioinsecticides) towards larvae of certain mosquito species (Frankenhuyzen, 2017) and references herein). In this study, we investigated the physiological and cellular mechanisms disturbed by *Btk* products which are responsible for developmental delay and growth defect. To best mimic environmental conditions, we used doses potentially present in the field after agricultural spraying. The lower dose we used corresponds to the amount deposited on the vegetables after one treatment. The highest dose reflects the amount that may be encountered after multiple treatments, even though data regarding *Bt* accumulation in the fields remain scarce. Indeed, the half-life of the entomopathogenic Cry1 toxins being estimated at ten days in the field (Hung et al., 2016), weekly sprays are recommended by the manufacturer and up to 10 are authorized. Regarding the persistence of spores, it has been shown that 72 hours after spraying of chards with *Btk* bioinsecticides, almost all of the *Btk* was still present at the surface of the leaves (Bizzarri and Bishop, 2008). Ten days later, the amount of *Btk* bioinsecticides corresponded to 50% of the initial dose sprayed on the leaves. The amount of *Bt* found on the leaves approaches normal values only 28 days after treatment (<10^1^ CFU/g) (Raymond et al., 2010). Furthermore, once *Bt* has reached the soil, it persists more than a year in the treated soil (Vettori et al., 2003). Our study highlights that such a persistence and accumulation of *Btk* bioinsecticides may affect the development of non-target insects. Nonetheless, the impacts are mainly due to the synergy between *Btk* spores and crystals since either the spores (*Btk*^*ΔCry*^) or the purified crystals alone do not promote detectable developmental delay or growth defects in *Drosophila* larvae. These data are in agreement with the study analyzing the safety of *Btk* purified Cry toxins on *Drosophila* larval development (Haller et al., 2017).

Interestingly, the effects we have characterized here are reminiscent of those observed on the target organisms *Helicoverpa armigera* and *H. zea*, *Spodoptera frugiperda* and *Choristoneura fumiferana*. Indeed, when fed with sublethal doses of *Btk* spores, all these target organisms display developmental delays and/or reduced pupal size (Bauce et al., 2006; Polanczyk and Alves, 2005; Rabinovitch et al., 2017; Sedaratian et al., 2013). Moreover, recent work has also shown that the potent pathogens *P. entomophila* and *P. aeruginosa* or huge amounts of the opportunistic bacterium *Erwinia carotovora carotovora* induce developmental delay and growth defects in *Drosophila* larvae (Houtz et al., 2019). These data suggest that developmental delay and/or growth defects must be general consequences of intestinal infections. The compromised gut digestive capacity of the infected larvae (whatever the insect) is likely the main reason for such developmental defects. Indeed, we and others have shown in the adult (Buchon et al., 2009; Chakrabarti et al., 2012; Erkosar et al., 2015; Loudhaief et al., 2017; Wong et al., 2016) or in larva (this work) that allochthonous bacteria induce a more or less widespread apoptosis of enterocytes leading to a decreased digestive and/or absorptive capacities. The developmental delay and the growth defect likely reflect the degree of damages suffered by the intestine. Houtz and colleagues observed a delay in pupation of up to 4 days using strong pathogens (Houtz et al., 2019) while we observed here a delay between 16 hours (Delfin) and 24 hours (Dipel) with *Btk* products. Therefore, while *Btk* behaves as a strong pathogen on target organisms, ultimately leading to death (in fact the expected effect), it has milder but significant impacts on the development of non-target insect. Our work demonstrates that the composition of the diet is important to overcome *Btk* intestinal infection. Increasing the protein content of the nutritive medium rescues both growth defects and developmental delay. Our data are in agreement with previous observations in mammals and insects infected with various pathogenic bacteria or viruses showing that protein intake is important to withstand attacks by pathogens, to support an efficient immune defense and to promote growth, while intake of carbohydrates is deleterious for all these functions (Bing et al., 2018; Lee et al., 2006; Reis, 2016; Rodrigues et al., 2015; Shikano and Cory, 2014; Soultoukis and Partridge, 2016). Moreover, our data show that the commensal bacterium *L. plantarum* also participates in the protection of larvae from *Btk* intestinal aggression in accordance with previous works involving commensal bacteria in gut protection against pathogens (Ubeda et al., 2017; Van Arnam et al., 2018). In *Drosophila*, *L. plantarum* has been shown to protect adult flies from *Sm-* or *P. aeruginosa*-induced mortality and this protection is specific since the commensal bacteria *Enterococcus faecalis* or *B. subtilis* do not operate as such (Blum et al., 2013). Our work strongly suggests that the benefit brought by *L plantarum* to the larvae of enduring *Btk* aggression passes through an enhanced protein digestion and/or absorption by the gut in line with previous observations (Erkosar et al., 2015). Finally, we did not observe a decrease in food intake suggesting that spores and commercial products do not induce avoidance behavior. Together, our data support the model that the impact of the *Btk* products on the development of the non-target organism *D. melanogaster* (and probably many other arthropods) depends on both the nutritional environment and the performance of its commensal flora. Consequently, spraying *Btk* products when environmental conditions are stressful would amplify the adverse effects of *Btk* products. Our work also sheds light on the pathogenic opportunism of *Btk* on a broad range of organisms owing to the conservation of the digestive tract architecture and functions (Pasco et al., 2015). For instance, the impacts of *Btk* on animals or humans suffering from dietary deficiencies (very often associated with dysbiosis) may be more deleterious than on individuals with a well-balanced diet. Of note, bacteria of the *B. cereus* group are well-known opportunistic pathogens responsible for foodborne poisoning (EFSA BIOHAZ, 2016; Wells-Bennik et al., 2016). Therefore, a more accurate monitoring of the presence of *Btk* in food and environment is needed to avoid any setbacks, especially in the current context where organic farming is gaining market share.

Finally, our investigations have unraveled two concomitant mechanisms allowing *Drosophila* larvae to overcome an intestinal infection. It was thought so far that the *Drosophila* larval gut was unable to regenerate due to the short period covering the larval stages (about 5-6 days) and the inability to give birth to news cells to replenish the damaged ones (Miguel-Aliaga et al., 2018). Only two choices were thus offered to the larvae: *Live or let die*. However, we have uncovered a third choice for the infected larvae: *Die another day*. First, we demonstrate that the adult midgut precursors (AMPs) can be diverted from their original fate. Indeed, upon aggression of the larval intestinal epithelium, the AMPs can engage toward a process of differentiation to give birth to enterocyte-like cells, which are intercalated in between AMP islets and enterocytes. We can assume that such newborn cells could plug the holes caused by the death of enterocytes, thus avoiding the entry of luminal bacteria into the internal milieu. Unfortunately, newborn cells do not fully differentiate and probably do not participate in digestive functions as efficiently as well-differentiated enterocytes. The addition of these misdifferentiated cells to dying enterocytes can explain the reduced protease activity of the intestine of infected larvae and the subsequent developmental delay and growth defect. The huge increase in the proliferation of AMPs that we observed at the end of L3 upon *Btk* spore ingestion could be explained by the necessity to replenish the right number of AMPs before pupation to achieve a complete adult gut metamorphosis and therefore produce a viable adult fly. These AMPs replenishment could also participate to the developmental delay we observed. The work very recently published by Houtz and colleagues reaches to similar conclusions (Houtz et al., 2019).

We have also uncovered a second unexpected mechanism involved in the maintenance of gut integrity: the change in enterocyte shape. So far, it has been shown in many species (insects, mammals or plants) that the loss of cells in response to stressors can be compensated by the rapid increase in the size of neighboring cells thanks to successive endocycles (Edgar et al., 2014). Here, we evidence an increase in enterocyte area upon *Btk* spore ingestion but concomitantly we have observed a thinning of the epithelium. Above all, we did not observe an increase in the ploidy of the enterocyte nuclei. Interestingly, it has been shown that old enterocytes in adult *Drosophila* midgut cannot re-enter the endocycle (Xiang et al., 2017). Therefore, as for old adult enterocytes, L3 enterocytes (born during embryogenesis 7 days earlier) are probably unable to re-enter a cycle of endoreplication to promote cell growth. Hence, changing their shape through flattening allows to increase the area covered by one enterocyte and to participate in maintaining epithelium sealing. Here again, the absence of endoreplication likely participates to the reduced digestive capacities of the intestine and therefore to the developmental delay and growth defects.

It has been proposed that lepidopteran larvae succumb to *Btk* spores and toxins because they are unable to maintain their gut integrity. Conversely, one mean for a lepidopteran larva to acquire resistance to *Btk* bioinsecticides may rely on its capacity to maintain/regenerate its intestinal integrity (Castagnola and Jurat-Fuentes, 2016; Castro et al., 2019; Dubovskiy et al., 2016). Non-target organisms are likely as such because they are able to regenerate their gut lining faster than *Btk* bioinsecticides destroy it. However, increasing the amount of *Btk* products provided to *Drosophila* larvae ultimately leads to their death (Babin et al., 2019), suggesting that the harmfulness of *Btk* products relies on both the amounts of *Btk* products ingested and the affinity of Cry toxins for more or less specific receptors. Altogether, these data suggest that *Btk* spores used as bioinsecticides may be not as safe as expected on non-target organisms. Moreover, the opportunistic virulence of *Btk* for many organisms should not be overlooked, especially for those suffering from nutritional stress, dysbiosis or with a compromised innate immunity. A situation that more and more species will encounter with the climatic changes occurring worldwide. Many investigations remain to be done to improve the use and efficiency of *Btk* products and to make their use as safe as possible for both handlers and consumers (wildlife or humans).

## MATERIALS AND METHODS

### Fly Stocks and genetics

Canton S (WT) (Bloomington #64349); *w, dlg::GFP cc01936* (FlyTrap) was provided by A. Spradling; *w; myo1A-Gal4* was a gift from N. Tapon; *w, UAS-GC3Ai*^*G7S*^ were kindly provided by Magali Suzanne and described in (Schott et al., 2017). Apoptosis was monitored on F1 progeny *w/w; myo1A-Gal4/+; UAS-GC3Ai*^*G7S*^/+ L3 larvae. The ReDDM fly stock to trace cell progenies in the midgut was kindly provided by Maria Dominguez (Antonello et al., 2015). To perform lineage tracing, synchronized eggs (*w/+; esg-Gal4 UAS-GFP/+; tub-Gal80*^*ts*^ *UAS-H2B::RFP/+*) were hatched on contaminated food (see below for the method) and maintained for 24h at 25°C. Then, L1 larvae were transferred and maintained at 29°C to alleviate the inhibitory effect of Gal80^ts^ on the Gal4 transcription factor. Intestine were dissected and observed at mid L3 (between days 4 and 5).

### Bacterial strains

*L. plantarum*^*WJL*^ that has been described in (Ryu et al., 2008) and sequenced in (Martino et al., 2015) was kindly provided by Bernard Charroux (IDBM, Marseille, France) and by Renata Matos (F. Leulier’s Lab, IGFL, Lyon, France). *Pseudomonas entomophila* (Vodovar et al., 2005) was kindly provided by Bruno Lemaitre’s Lab (EPFL, Lausanne, Switzerland). *P*_*e*_ and *L. plantarum*^*WJL*^ were grown as described in (Loudhaief et al., 2017). The *Btk*^*ΔCry*^ strain (identified under the code 4D22) (Gonzalez et al., 1982) was obtained from the *Bacillus* Genetic Stock Center (www.bgsc.org). Delfin and Dipel formulated bioinsecticides were bought in an agricultural shop center. The strain *Btk* ABTS-351 was isolated from Dipel on LB agar.

### Spore preparation

*Btk* vegetative cells were let to grow and sporulate during two weeks at 30°C with shaking in 2L of PGSM medium. The culture was then heated 1h at 70°C to eliminate vegetative cells and centrifuged for 15 min at 5000g. The pellet was first washed with 1 L of 0.15M NaCl and twice with sterile water. The final pellet was suspended in 30 mL of sterile water, aliquoted and lyophilized 24h-48h. The spore titration was determined by serial dilution of a known weight of lyophilized spores and counting the number of “colony forming unit” (CFU) on LB agar after incubation at 30°C for 18h.

### Crystal purification

Two grams of the commercial product were suspended in 60 mL of sterile water, dissolved for 5h at 4°C with agitation and then sonicated 4 times 15s with a frequency set at 50% (Fisherbrand™ Model 505 Sonic Dismembrator). Ten milliliters of the homogenate were deposited on a step density gradient of 67/72/79% sucrose and centrifuged at 100 000g for 18h at 4°C. Crystals were collected on the interfaces between the different sucrose phases and washed twice with sterile water. The final pellet was suspended in 12 mL of sterile water, aliquoted and lyophilized 24h-48h.

### Diet composition

*Drosophila* were reared at 25°C on conventional medium (referred as the “poor-protein” medium in this study): 8% cornmeal (AB Celnat), 2% yeast (Springaline® BA10/0–PW, Amcan), 2.5% sucrose, 0.8% agar (ref 20768-361, VWR) and 0.6% methylparaben (Tegosept,, ref 789063 Dutscher). The protein rich medium contains 10% yeast instead of 2%.

### Intestinal CFU counting

Colony forming unit counting were described in (Loudhaief et al., 2017).

### Commensal flora estimation

Ten flies or ten mid-third instar larvae were washed first with ethanol 70% for 30 seconds, then washed with PBS and crushed in 500 μL of LB medium with a Tissue Lyser (Qiagen ref. 85600) at 50Hz for 5 minutes. The homogenate was plated on different selective media to identify the common commensal bacteria. MRS agar (Sigma ref 69964 + 1 mL of Tween 20/L) at 37°C in anaerobic was used to identify *Lactobacillus plantarum* and *brevis*. BHI agar (Sigma ref 70138) at 37°C allowed the identification of *Enterococcus faecalis*. LB agar (Fisher ref BP1426-2) at 37°C allowed the identification of *Escherichia coli* and *Staphylococcus aureus*. Mannitol medium (Bacto peptone 3g/L, Yeast extract 5g/L, D-Mannitol 25g/L, agar 15g/L) at 30°C allowed the identification of *Commensalibacter intestini* and *Acetobacter pomorum*.

### Axenic flies

Adult Canton S flies were let to oviposit eggs for two days at 25°C and 12h/12h day/night cycle on our conventional medium supplemented with a cocktail of four antibiotics: ampicillin, kanamycin, tetracycline and erythromycin (50 μg/mL each). On the next generation, newly hatched adults were immediately removed and transferred on fresh conventional medium containing the four antibiotics. This operation was repeated until flies with axenic gut were obtained. The presence of commensal flora was checked at each fly generation (Fig. S1). Axenic flies were obtained after 6 generations.

### Oral infection of larvae

Adult flies were let to oviposit eggs for 4h at 25°C on laying nests containing grape juice medium (25% grape juice, 3% agar, 1.2% sucrose, 2% ethanol, 1% acetic acid). Twenty eggs were collected and transferred in a *Drosophila* vial containing 2g of the desired medium mixed with *Btk* spores, *Btk* crystals or the naphthalene additive. The conditions used for 2g of medium and 20 eggs were: 10^7^ CFU of *Btk* spores for the 1X dose; 10^8^ CFU of *Btk* spores for the 10X dose; 0.67mg of *Btk* purified crystals (equivalent to the 10X dose and considering that crystals represents 15-30% of spore weight (Agaisse and Lereclus, 1995; Monro, 1961; Murty et al., 1994)); 10^8^ CFU of *L. plantarum* or *Pe*; 0.4mg of naphtalene-2-sulfonic acid sodium salt (ref A13066, Alfa Aesar) (equivalent to the 10X dose). Vials were incubated at 25°C with a 12h/12h day/night cycle until further processing.

### Pupation curve

Immobilized larvae forming white pupae were counted. The number of pupae was expressed as the percentage of the final number of pupae. While 20 eggs were deposited in each tube, in some conditions, especially with Dipel and Delfin, we never got 20 pupae probably due to previously described larval lethality induced by the *Btk* products (Babin et al., 2019). Table 1 summarizes the developmental time at which 10%, 50% and 90% of the larvae were immobilized for all the conditions we tested. For each condition described above, the experiment was performed at least in three experimental blocks of triplicates.

### Pupal size

On day 7, 8 and 9 after egg laying, pupae were gently removed from the vial, washed in water and slightly dried with tissue paper. Pupae were placed on a microscope slide and pictures were taken with a numeric microscope VHX-2000 (Keyence). Pupal length and width were measured with Image J software and pupal volume was calculated using the following formula for ellipsoid: V= [(4/3) × 3,14 × (L/2) × (l/2)^2^] where V is the volume, L the length and l the width of the pupa.

### Larval feeding behavior and Food intake

Quantification of larval food intake was performed according to (Bjordal et al., 2014) with the following modifications: we deposited a mix of 50% yeast/0.75% of the food blue dye Erioglaucin (ref 861146, Sigma) in the center of a Petri dish lid (diameter: 35mm) to feed 20 mid-third larvae (previously infected or not) *ad libitum*. After 2h at 25°C, larvae were collected and washed with water. The number of white larvae (indicating no feeding) and blue larvae was counted. Ten blue larvae were transferred into a microtube containing 500 μL of PBS, crushed at 50Hz for 2 minutes with a TissueLyser (Qiagen, ref 85600) and centrifuged at 10000g for 5 minutes at 4°C. The amount of Erioglaucin in the supernatant was quantified at 629 nm with a spectrophotometer (SpectraMax® Plus384, Molecular device).

### Midgut permeability

The same technique as described above was used except that the mix contained 50% yeast and 1.25 mg/ml Dextran FITC (ref 46944, Sigma) in water. Twenty previously orally infected mid-third instar larvae were placed in the mix. After 2h at 25°C, the larvae were collected, washed with water and placed at least 30 minutes at 4°C. Pictures were taken with a fluorescent stereoscopic AxioZoom V16 (Zeiss) microscope.

### Metabolic assay

Intestinal protease activity assay was performed as described in (Nawrot-Esposito et al., 2017) with 8 mid-third instar larval midguts per condition. Lipids were assayed as described in (Tennessen et al., 2014). The tri- and diacylglycerol (TAG and DAG) were measured on 3μl of hemolymph obtained from 5 days old larvae using the Triglyceride Reagent (ref T2449, Sigma) and Free Glycerol Reagent (ref F6428, Sigma). The metabolite quantity was reported per mg of protein or per μL of hemolymph. Protein assay was achieved with Biorad Protein Assay Dye Reagent Concentrate (ref 5000006, Biorad). All experiments were performed at least 3 times in triplicates.

### Lipids labelling

Five days after infection (mid-third instar larvae), ten larval guts were dissected in PBS and fixed 40 min in 4% formaldehyde in PBS. Guts were then washed twice in PBS and further incubated for 30 min. either in 5μg/ml of BODIPY™ 493/503 (ref D3922, Invitrogen™ Molecular Probes™) or in a 0.06% Oil-red-O solution (ref 189400250, Acros Organics). Guts were then washed twice with PBS. For BODIPY™ labelling guts were mounted in Fluoroshield/DAPI (ref F6057, Sigma) and immediately observed under a fluorescent microscope (Zeiss Axioplan Z1 with Apotome 2). For Oil-red-O labelling, guts were mounted in 80% Glycerol/PBS and observed with a numeric microscope (VHX 2000, Keyence).

### Immunohistochemistry, image capture and processing

Ten guts of late third instar larvae were dissected in PBS and fixed for 40 min. in 4% formaldehyde in PBS. Guts were then washed twice with PBS-0.1% Triton X100, incubated for 3h in Blotto (2.5% FBS, 0.1% Triton X-100, 0.02% sodium azide in PBS) and further incubated with the primary antibody overnight at 4°C in Blotto. The following dilutions were used: Mouse anti-Armadillo (DSHB #N2 7A1) at 1/50; Rabbit anti-Phospho-histone-3 (ref 9701S, Cell signaling) at 1/500. Guts were washed once with Blotto before incubation in the secondary antibody and/or phalloidin (2h at room temperature in Blotto). The secondary antibodies were the followings: AlexaFluor™ 546 donkey anti-Rabbit IgG (ref A10040, Invitrogen) used at 1:500; AlexaFluor™ 647 goat anti-Mouse IgG (H+L) (ref A21235, Invitrogen) used at 1/500; Alexa Fluor™ 555 Phalloidin (ref A34055, Invitrogen) used at 1/1000. Guts were finally washed several times in PBS, mounted in Fluoroshield/DAPI (ref F6057, Sigma) and observed with a Zeiss Axioplan Z1 with Apotome 2 microscope. Images were analyzed using ZEN (Zeiss) software or Photoshop. For cell death monitoring (in *myo1A-Gal4 UAS-UAS-GC3Ai*^*G7S*^ *larvae*), guts were dissected in PBS, immediately mounted in 80% Schneider medium/20% glycerol and pictures were rapidly taken with a fluorescent stereoscopic microscope (AxioZoom V16, Zeiss). Image acquisition was performed at the Microscopy platform of the Institut Sophia Agrobiotech (INRAE 1355-UCA-CNRS 7254-Sophia Antipolis).

### Polyploidy estimation

Ploidy was calculated by measuring the area of the nuclei of late L3 larvae. Only round nuclei were analyzed. The average nucleus area of diploid adult midgut precursor cells was calculated. This average area (8μm²) was assigned to the 2C ploidy (C corresponding to chromatin copy number). The theoretical average area of 4C, 8C, 16C, 32C and 64C nuclei was calculated on the basis of the measured 2C average area. 109 nuclei for the control larvae, 184 nuclei for *Btk*^*ΔCry*^ 10X larvae and 306 nuclei for Dipel 10X larvae were counted and binned into ploidy groups.

### Statistics

When “n” was equal or superior to 30, statistical analysis was performed using a parametric T-test. An F-test was systematically done before applying the T-test to verify the homogeneity of variances. When “n” was inferior to 30, we used the non-parametric pairwise comparisons of the Wilcoxon-Mann-Whitney test. Significance symbols are the following: *** (P<=0.001); ** (P<=0.01), * (P<=0.05).

## Supporting information

SupFigures

## Acknowledgement

We would like to thank Nathalie Arquier and Rénald Delanoue for their help in food intake measurement, lipid dosage and staining, and pupae volume measurement. We thank Julien Colombani for the fruitful discussions, Pierre Barbero for discussions and correction of the manuscript, Bernard Charroux, François Leulier and Renata Matos for providing *L. plantarum*. We also thank Marine Chesneau for technical help during her internship. Thanks to Olivier Pierre from the Microscopy platform (Institut Sophia Agrobiotech, INRAE 1355-UCA-CNRS 7254-Sophia Antipolis) for help with the newly acquired Zeiss AxioZoom Microscope and to Raphaël Rousset for help with statistical analyses and manuscript correction. Finally, we warmly thank Rénald Delanoue for his help in designing the experiments and correcting the manuscript.

## Author contributions

**Marie-Paule Nawrot-Esposito:** Methodology, Validation, Formal analysis, Investigation, Resources, Data Curation, Writing - Original Draft **Aurélie Babin:** Methodology, Validation, Formal analysis, Investigation, Writing - Review & Editing **Matthieu Pasco:** Investigation **Marylène Poirié:** Writing - Review & Editing, Funding acquisition **Jean-Luc Gatti:** Validation, Writing - Review & Editing, Funding acquisition **Armel Gallet:** Conceptualization, Validation, Formal analysis, Data Curation, Writing - Original Draft, Writing - Review & Editing, Visualization, Supervision, Project administration, Funding acquisition

