## Supplementary figures and images for "*Bacillus thuringiensis* bioinsecticides induce developmental defects in non-target *Drosophila melanogaster* larvae"

### SupFigures

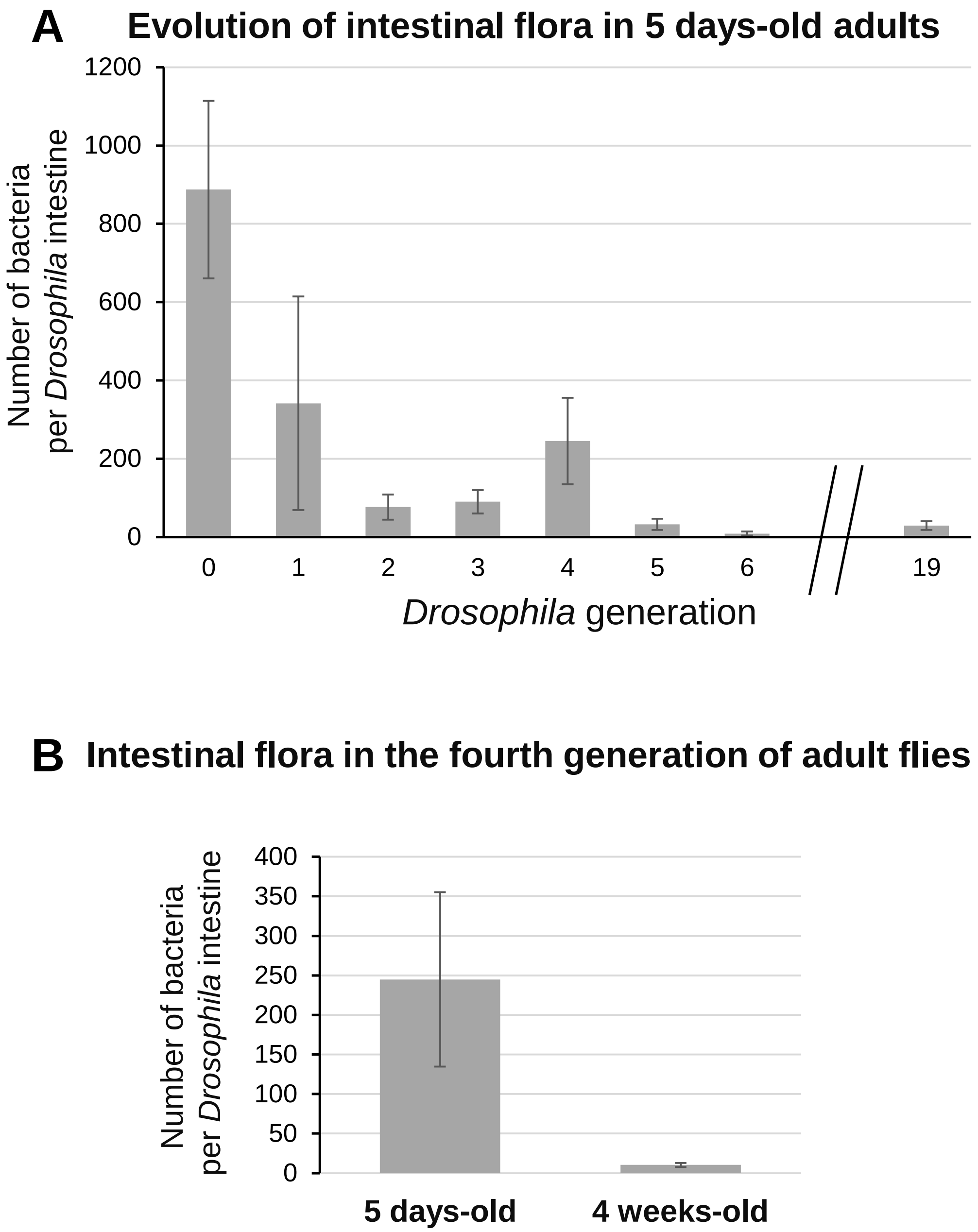

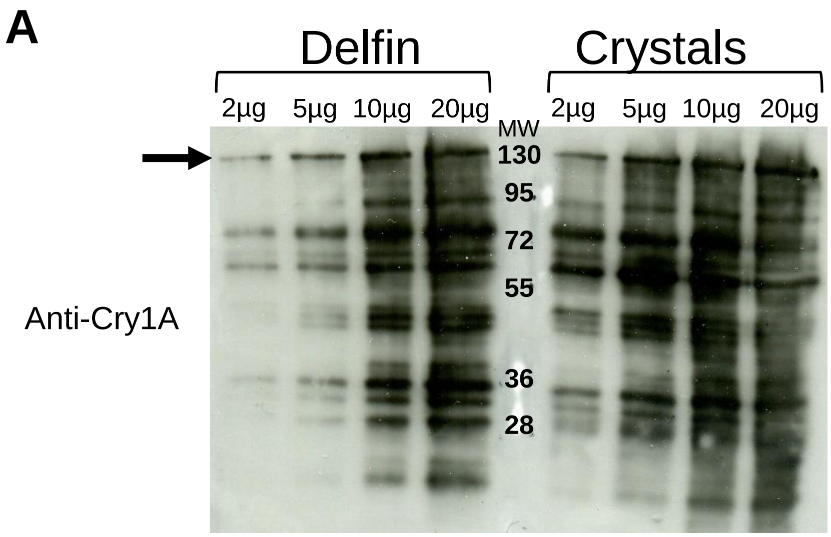

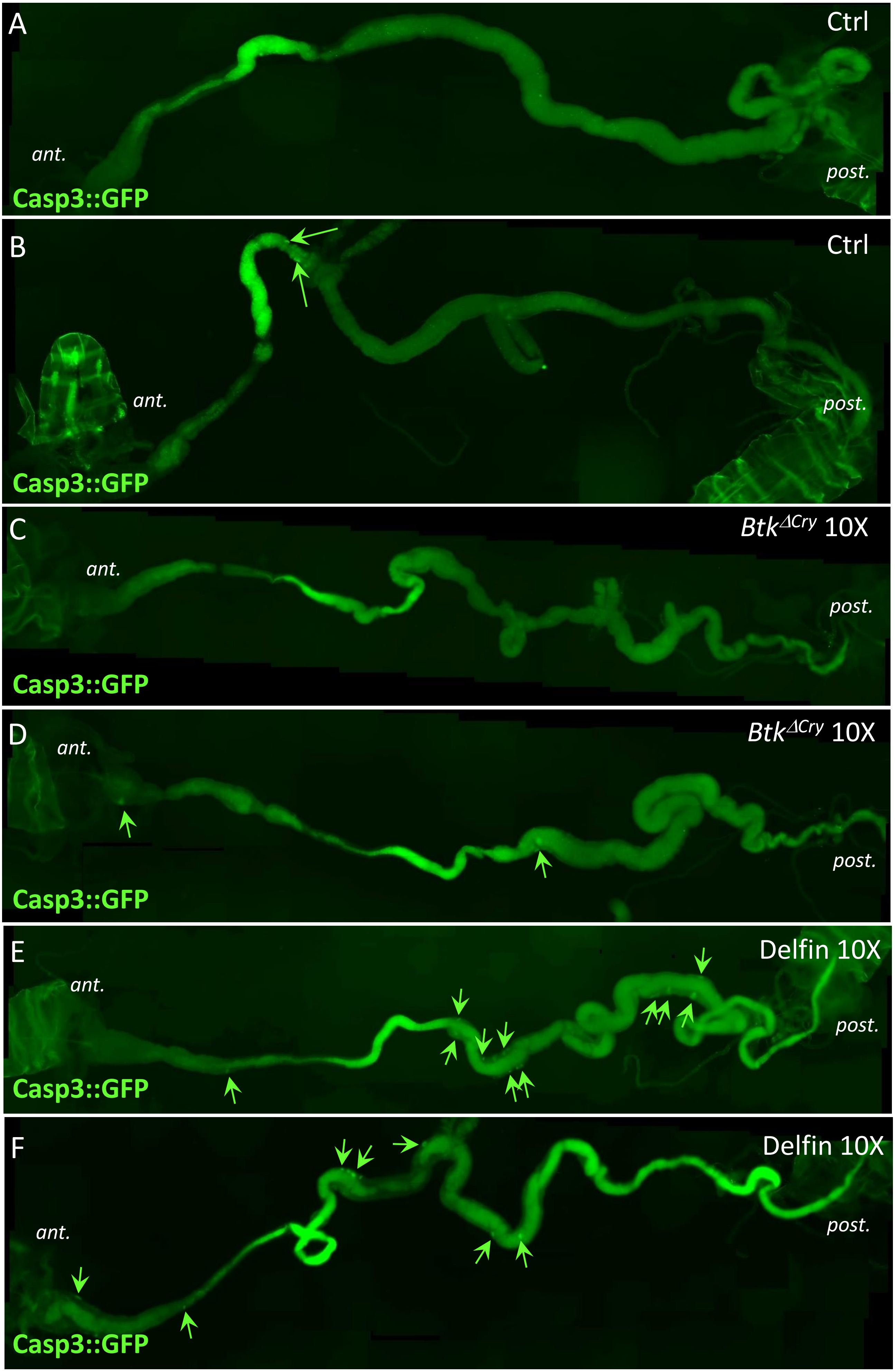
