## Supplementary material for "*Bacillus thuringiensis* bioinsecticides induce developmental defects in non-target *Drosophila melanogaster* larvae": SupLegends

**Supfigure 1: Commensal flora of Canton S adult *Drosophila melanogaster***

**(A)** Monitoring of whole intestinal commensal flora over 19 generations during the generation of the axenic strain. We considered that axenic flies were obtained at the $5^{th}$ generation. We used axenic the flies from the $7^{th}$ generation onwards to perform our experiments. **(B)** Intestinal commensal flora of young (5 days old) and aged (4 weeks old) adult flies of the $4^{th}$ generation during the axenic establishment procedure. Error bars represent SEM.

**Supfigure 2: Evaluation of the crystal amount**

Western Blot (WB) using a Rabbit antiCry1A antibody (1/5000, Babin et al., 2019). Equal amounts (2, 5, 10 and 20µg) of Delfin (left part of the WB) and purified crystal (right part of the WB) were deposited on an SDS PAGE. We estimated that Crystals represent between 25% and 30% of the Delfin weight. The band at 130kDa corresponds to the Cry1A protoxins (upper arrow). MW: Molecular Weight in kiloDalton (kDa).

**Supfigure 3: Caspase 3 activity in the L3 midgut**

**(A-F)** *myo1A>Casp::GFP* larvae raised on control a medium (A nd B) or on a medium contaminated with $Btk^{\Delta Cry}$ 10X (C and D) or Delfin 10X (E and F). Arrows point enterocytes in which Caspase 3 is activated. Anterior is toward the left.
